# Chemodiversity affects preference for *Tanacetum vulgare* chemotypes in two aphid species

**DOI:** 10.1101/2023.03.16.532937

**Authors:** Annika Neuhaus-Harr, Lina Ojeda-Prieto, Elisabeth Eilers, Caroline Müller, Wolfgang W. Weisser, Robin Heinen

**Affiliations:** Technical University of Munich, Department of Life Science Systems, School of Life Sciences, D-85354 Freising; Department of Chemical Ecology, Bielefeld University, 33615 Bielefeld, Germany; CTL GmbH Bielefeld, Krackser Str. 12, 33659 Bielefeld, Germany

**Keywords:** aphids, attractiveness, choice assays, genotypic variation, intraspecific phytochemical diversity, plant diversity, terpenes, terpenoids

## Abstract

Plants of the same species often strongly differ in morphological traits, as well as in the abundance and composition of specialized metabolite profiles. Specialized metabolites can act as mediators of interactions on plants, and affect insect presence and abundance in the field. However, how specialized chemistry shapes plant attractiveness to herbivorous insects is not fully understood. Here we used common tansy (*Tanacetum vulgare* L., Asteraceae) – a perennial plant that is highly diverse in terpenoid composition and is known to have variable chemotypes – to test whether 1) plants with different chemotype profiles differ in attractiveness to two specialized aphids, *Macrosiphoniella tanacetaria* and *Uroleucon tanaceti*, in pairwise choice assays. Furthermore, we tested whether 2) the diversity of the terpenoid blend affects aphid attractiveness. Lastly, we tested how 3) plant chemical traits relate to plant morphological traits, and which best explain aphid preference. We found that *M. tanacetaria* preferred two out of five chemotypes, dominated by α-thujone/β-thujone and β-trans-chrysanthenyl acetate, respectively, while avoiding a chemotype dominated by α-pinene/sabinene. *U. tanaceti* showed no clear preference towards chemotypes, but when given the choice between chemotypes dominated by α-thujone/β-thujone and by α-pinene/sabinene, they preferred the former. Importantly, plant attractiveness to aphids tended to be negatively correlated with chemodiversity, i.e., the number of terpenoid compounds, in *M. tanacetaria*, but not in *U. tanaceti*. Interestingly, the approximate concentration and number of terpenoid compounds was generally higher in larger and bushier plants. Hence, we did not observe a trade-off between plant growth and defence. We conclude that plant chemical composition affects plant attractiveness to aphids and hence may contribute to variation in natural aphid colonization patterns on plants of the same species.

## Introduction

Understanding relationships between plants and herbivores is an important goal in ecology. How variation in plant diversity shapes herbivory has been a subject of study for many decades (Scherber et al. 2010, Weisser et al. 2017). For a long time, between-species diversity was believed to be more important than within-species diversity as a driver of ecosystem processes (Des Roches et al. 2018). Intraspecific diversity - which includes the variation between individuals, and richness and abundance of genotypes and phenotypes within a population - has only recently gained attention as an important driver of ecological processes (Raffard et al. 2019). It has been shown that intraspecific variation in plant geno- and phenotypes can structure food webs, such as arthropod-plant interactions (Crutsinger et al. 2006, Poelman et al. 2008, Bálint et al. 2016). As plants of the same species can differ strongly in various traits, understanding how this variation contributes to shaping interactions between plants and interaction partners is currently an important goal in plant ecology.

One important dimension of intraspecific variation in plants is chemodiversity; the variation in chemical composition between individuals of plants of the same species (Wetzel & Whitehead 2020). Primary metabolites are important as the building blocks of plants, and these compounds are comparatively similar across the plant kingdom (Weng, Philippe & Noel 2012). Specialized metabolites, on the other hand, play critical roles as mediators of interactions and plants have evolved a staggering inter- and intraspecific diversity in these specialized compounds (Pichersky & Gang 2000; Iason, Dicke & Hartley 2012). Such chemical compounds can either be stored, emitted, or emitted after a stress event (Clancy et al. 2016; Holopainen and Gershenzon 2010). Intraspecific variation in specialized metabolite profiles is known to affect the structure of associated herbivore communities including phloem-feeding insects such as aphids (Schmitz 2008, Poelman et al. 2008, Richards et al. 2015, Bálint et al. 2016, Bustos-Segura et al. 2017, Salazar et al. 2018, Volf et al. 2019, Singh et al. 2021, Whitehead et al. 2021), but few studies have shown how chemodiversity structures herbivore abundance and community structure in natural conditions, by affecting herbivore preference, and bottom-up and top-down (predation) processes.

Chemodiversity may mediate presence and abundance of insect herbivores on a plant in the field, which likely results as a combination of i) attractiveness of the plant for the insect herbivore, ii) plant quality (i.e., bottom-up effects), and iii) the influence of higher trophic levels on the insect herbivore through predation (i.e., top-down effects). Many specialized compounds likely evolved to deter and repel herbivores (Herms & Mattson 1992, Kessler & Baldwin 2001, Whitehead et al. 2021), but particularly specialized herbivore species may use particular compounds as host-finding cues (Nishida 2014, Wink 2018). As such, insects can be repelled or attracted, particularly by emitted volatile organic compounds, or VOCs (Clancy et al. 2016, Jakobs & Müller 2019). Once an insect herbivore has arrived on the plant, stored and emitted compounds can affect herbivore performance, for instance by feeding deterrence, or influencing metabolic processes, but these processes largely depend on metabolite-specific modes of action (Unsicker et al. 2009, McCormick et al. 2012, Jakobs, Schweiger & Müller 2019). Emission of VOCs can also attract natural enemies of herbivores, thereby acting as an indirect layer of defense for the plant via top-down regulation (Dicke 2009, Unsicker et al. 2009, McCormick et al. 2012, Rzanny et al. 2013). As these three processes act on plant individuals simultaneously, their independent weights cannot reliably be inferred from natural colonization data collected from the field. It is therefore important to understand how intraspecific chemical variation mediates processes driving herbivore preference and performance in a controlled environment.

Besides intraspecific variation in chemical traits, individual plants of the same species may also differ strongly in morphological traits. Various studies point out that traits related to growth or to structural defenses may play an important role in driving interactions between plants and insects (Herms & Mattson 1992; Agrawal & Fischbein 2006). For instance, in a study on wheat plants, Batyrshina et al. (2020) found higher numbers of aphids on fast-maturing than on slow-maturing wheat plants, and trade-offs between plant growth and defence are thought to be common in nature (Coley et al. 1985; Herms & Mattson 1992). Furthermore, plants with stronger mechanical defences tend to be better defended against insect herbivory (Caldwell, Read & Samson 2015). It is plausible that chemical traits are linked to morphological traits, together driving insect preference. For instance, in common tansy, (*Tanacetum vulgare* L., Asteraceae), plants with a higher storage of camphor were found to have taller shoots than those with lower amounts, while plants containing davadone- D or artemisia ketone developed more flower heads, taller corymbs, and delayed flowering compared to plants with a lower content of these compounds (Keskitalo et al. 2001). Furthermore, *T. vulgare* from different origins (e.g., North America and Europe) have been found to differ in both morphological and chemical traits, and exhibit negative correlations between reproductive biomass and terpene concentrations (Wolf et al. 2011). Moreover, Hayashi, Tahara and Ohgushi (2005) found that pubescence density and leaf mass per area differed between the two chemotypes of *Salix sachalinensis*. The interplay of morphological and chemical plant traits are plausible drivers of insect communities, but their relative contribution as drivers is not fully understood.

*Tanacetum vulgare* is a perennial, aromatic plant that has a large geographic distribution and is associated with a complex herbivore community including mono-, oligo- and polyphagous aphids (Schmitz 1998, Keskitalo et al. 2001, Kleine & Müller 2011). Tansy is rich in different mono- and sesquiterpenoids and plants can be divided into chemotype groups based on the composition of terpenoids (Keskitalo et al. 2001, Kleine & Müller 2011). Previous studies and breeding experiments showed that terpenoid composition has a genetic basis in tansy (Keskitalo et al. 2001). Specialized aphids are thought to be adapted to harmful metabolites in tansy and to use emitted volatiles for finding host plants (Schoonhoven et al. 2005, Jakobs & Müller 2019). Aphid colonization, growth rate and survival, and even the structure of aphid genotypes were found to be partly explained by tansy chemotypes under natural colonization (Senft et al. 2017, 2019, Zytynska et al. 2019, Clancy et al. 2018). Specialized aphids show preferences towards specific chemotypes (Jakobs & Müller 2018), but whether this is driven by chemotype composition or concentration of compounds when several chemotypes are on offer, still needs investigation.

In this study, we used six tansy chemotypes to investigate how differences between plant individuals in chemical traits shapes attractiveness to the specialized aphids *Macrosiphoniella tanacetaria* and *Uroleucon tanaceti* in factorial pairwise choice-assays under lab conditions. We hypothesized that 1) the two aphid species would show species-specific attraction to distinct tansy chemotypes, as previously shown by Jakobs and Müller (2018, 2019). Furthermore, we hypothesized that 2) chemodiversity negatively correlates with attractiveness, under the assumption that most specialized metabolites repel antagonistic organisms. We specifically investigated whether approximate concentrations of individual compounds predicted attractiveness. We also investigated how chemical composition relates to morphological traits. We hypothesized that 3) growth-related traits would trade-off with chemodiversity, under the assumption that maintaining chemodiversity is costly and limits available resources for growth. We specifically explored whether chemical or morphological traits are more important in driving insect preference towards specific plant chemotypes.

## Materials and methods

### Chemotypic characterization of tansy lines

In 2019, leaf and seed samples of 27 tansy plants (hereafter: mothers) were collected in Jena, Germany (50.93°N, 11.58°E), and chemotyped based on their terpenoid profiles. Terpenoids were analysed as in Kleine & Müller (2011) and Wolf et al. (2012). Slight modifications were made as in Eilers et al. (2021). The leaf material was freeze-dried, homogenised, weighed and extracted in heptane, adding 1-bromodecane as internal standard. Extracts were centrifuged and the supernatants analysed using gas chromatography coupled with mass spectrometry (GC-MS; GC 2010plus – MS QP2020, Shimadzu, Kyoto, Japan) on a semi-polar column (VF-5 MS, 30 m length, 0.2 mm ID, 10 m guard column, Varian, Lake Forest, United States) in electron impact ionisation mode at 70 eV, with helium as carrier gas. Samples were injected at 240 °C with a 1:10 split. A starting temperature of 50 °C was kept for 5 min, ramping up to 250 °C at 10 °C min^-1^, then increasing with 30 °C min^-1^ to a final temperature of 280 °C, hold for 5 min. An alkane standard mix (C7–C40, Sigma Aldrich, Taufkirchen, Germany) was measured regularly between samples. Terpenoids were identified based on their retention indices and by comparing spectra to various reference sources. Terpenoids were semi-quantified based on the peak area of the total ion chromatogram, related to sample dry mass and the internal standard.

Plant chemical profiles were clustered using unsupervised hierarchical k-means clustering with the hclust() function, performed on the dissimilarity matrix that was obtained from the approximate absolute terpenoid concentrations using the dist() function. Absolute values were used here, as attractiveness of a chemical profile is likely to be a combination of blend composition and approximate total concentration, the latter of which would be taken out by normalizing the terpenoid profiles. The number of clusters, *k*, was obtained using the elbow method. We selected a *k* = 7 for mothers, of which six were used for further chemotype selection. All statistical analyses were performed in R (R Core Team 2021).

Tansy is outcrossing, implying that daughter plants originating from seeds may have a degree of maternal similarity, but may also be quite different chemical profile because of paternal variation. For the preference experiment we aimed for a broad range of plants differing in chemotypic composition. Hence two mothers were randomly selected per cluster and the collected seeds were mass-sown in seedling trays in November 2020. Three weeks later, ten healthy seedlings were selected from each mother plant lineage and transplanted to 10-cm pots filled with standard potting substrate (Stender potting substrate C 700 coarse structure, 1 kg NPK minerals/m³, pH 5.5–6.0) and transferred to a greenhouse compartment at the greenhouse facilities at TUM Plant Technology Centre (with 16h:8h L:D and supplemental lighting that was turned off when outdoor light was > 40 klx). In total, this resulted in a set of 120 plants (hereafter: daughters). After seven weeks of growth in the greenhouse, the ultimate 3-4 leaflets from the youngest fully expanded pinnate leaf from the top of each plant were harvested. Samples were flash-frozen in liquid nitrogen, and subsequently freeze-dried for chemotyping. All daughters were grown under the same greenhouse conditions until July 2021, after which they were transplanted to a common garden on-site and watered well until fully established.

A second unsupervised hierarchical clustering analysis was then performed on the daughter terpenoid profiles. For the daughter profile clustering, the elbow *k* was at 5 - 7 clusters, which is why *k* = 6 clusters was used for daughters. Hierarchical k-means clustering of terpenoid profiles resulted in six large clusters, without any small outlier clusters driven by strongly deviating individual profiles (Fig. 1). Daughter lineages did not typically resemble mother lineages in their terpenoid profiles, and therefore the observed clusters did not reflect the mother lineages (Supplementary Fig S1). From each cluster, we selected three daughters from the same mother lineage. When more than three daughters were available from a mother in a cluster, we selected the three daughters clustering closest together. Our chemotype selection resulted in six maternal chemotype lines with three daughter replicates for each cluster (Fig. 1) that were used for preparing plants for the preference experiment.

**Figure 1:**
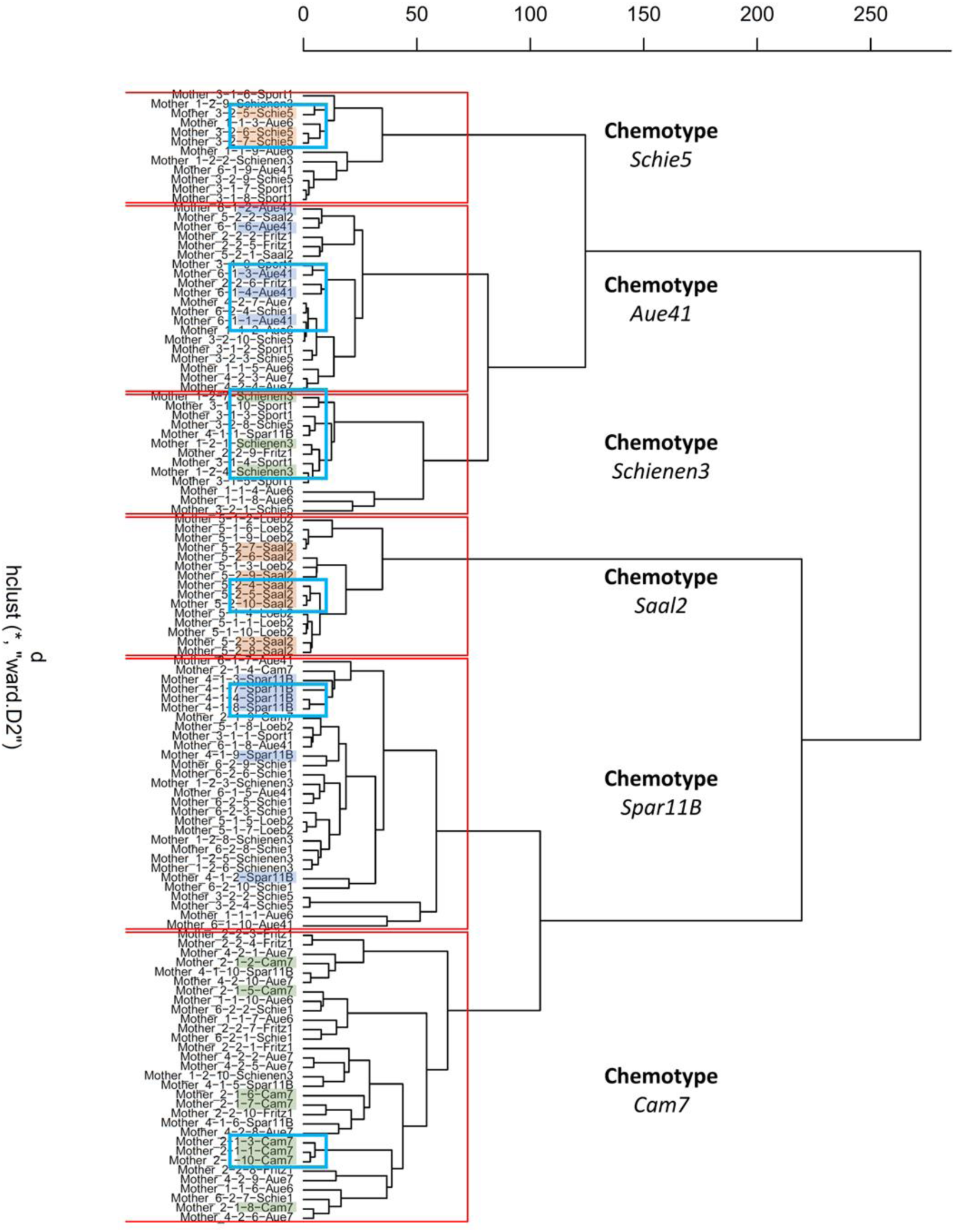
Hierarchical clustering tree for all 120 daughters terpenoid profiles. Clusters (k = 6) are indicated in red boxes. For the selection, in each cluster a different mother lineage was chosen, and three daughters were manually selected and highlighted. The final selection of daughters of each cluster are indicated in blue boxes.

### Propagation of plant material for aphid choice assays

In August 2021, fresh plant material was taken from the 18 selected daughters, and shoot cuttings were prepared by cutting stem parts 1-2 cm below and 4-5 cm above a leaf node. The leaf size was reduced by roughly 60%, to reduce evapotranspiration. The stem cuttings were pressed in seedling trays filled with potting substrate that was soaked. Cuttings were covered by a transparent plastic hood that was gradually opened after the plants established roots and showed first signs of shoot formation after three weeks. After acclimatizing the cuttings for two weeks, they were repotted to 10 cm pots, and later to 17 cm pots, to avoid pot limitation before their use in choice assays in December 2021. Plants were bottom-watered automatically and fertilized with Universol Blue fertilizer (18% N - 11% P - 18% K; ICL Deutschland, Nordhorn, Germany), target electrical conductivity of 1.0. No chemical insecticides or fungicides were used during plant propagation. Due to low propagation success of the Aue41 lineage, only the three daughters of the remaining five chemotypes were used in pair-wise aphid choice assays.

### Morphological traits

Morphological traits were measured non-destructively ten weeks after sowing for each of the 120 daughter plants in February 2021. The number of stems, leaves, and total number of nodes per plant were counted, height was recorded in cm, and internode length was derived by dividing height (in cm) by the number of nodes. From the youngest fully expanded leaf, the total number of leaflets from one leaf, leaf length, and leaflets density (number of leaflets / petiole length) were assessed, and chlorophyll content was measured using a chlorophyll meter (Konica Minolta SPAD-502Plus, Tokyo, Japan). The leaf was harvested and photographed in a flattened position with a reference mark for determination of leaf surface area using ImageJ®. The leaves were dried for 72 hours at 60°C to determine dry weight, specific leaf area, and specific leaf mass.

### Aphid rearing

Adults and nymphs of *Macrosiphoniella tanacetaria* and *Uroleucon tanaceti* were collected in August 2021 tansy plants in a field in Dürnast (48.4049230,11.6898638) and kept in plastic cages at room temperature with supplemental light at long day regimes (16h:8h L:D). Each aphid colony was provided with two to four tansy plants at a time. Plants for feeding the aphid colonies were collected from different locations in the vicinity of Technical University of Munich campus in Freising, Germany, and were unrelated to the used chemotypes for this study to avoid systemic bias in preferences in the aphid colonies. For each testing round, roughly 100 unwinged adult aphids of each species were placed on individual fresh plants. After 48 h, adult aphids were removed, and all nymphs kept on the plant. Subsequently, the cohorts were kept in a Fitotron® standard growth chamber 120 (21/16°C, 60% RH, Weiss Technik, Giessen, Germany) for eight days. After that the aphids were starved in a petri dish with a small piece of wet tissue for 24 h before the start of choice assays.

### Pairwise choice assays

Five out of the six *T. vulgare* maternal chemotypes with three replicate daughters were used for choice assays. *A priori*, a full-factorial series of pairwise choice assays was designed, in which aphids could choose between two different chemotypes (Suppl. Table S1). Within each replicate series, all chemotypes were tested against each other using randomly picked daughters. However, we ensured that all possible chemotype-daughter combinations were used in multiple replicate rounds. Choice assays were conducted in three rounds on three different days with different aphid cohorts that were standardized by age (9-10 days old), but within one week (9th - 16th December 2021). Multiple replicate rounds were performed in each experimental day. In total we performed 23 replicate rounds for *M. tanacetaria,* and 13 for *U. tanaceti*, as their cohorts numbers were substantially lower in number. *Tanacetum vulgare* daughters were assigned beforehand to ensure that each daughter was tested against each other daughter from the other chemotypes. However, as *M. tanacetaria* was used in five rounds which had been originally assigned to *U. tanaceti*, numbers of combinations of tansy daughters differed (for more details see Suppl. Table S1).

For the choice assay setup, the second and third youngest fully expanded leaf of a plant of the respective chemotype were selected, the first three leaflets discarded, and the following leaflets clipped with scissors. Plant leaflets were cut between 9:00 and 10:00 and placed in petri dishes. In petri dishes (14.5 cm diameter), two different leaflets were placed at equal distances from the centre, 8 cm apart from each other. The sides for each replicate alternated between replicate rounds to account for external bias. After preparing all the leaflets, one starved aphid was placed in the centre of the petri dish, and the petri dishes were sealed with Parafilm®, to prevent the leaves from drying quickly and placed in a TUM Model EcoSystem Analyser (TUM*mesa*) climate chamber. The LED lighting system in these chambers generates homogenous light conditions, aimed at reducing spatial effects. Petri dishes were placed onto a stainless-steel table and left for 24 h (16h:8h L:D photoperiod, 21°C, 60% RH). Aphid preference was recorded after two, five and 24 h. Each aphid and leaf were only used once. Dead aphids were excluded from further analyses.

### Statistical analysis of aphid preference

To test our first hypothesis, whether aphids show species-specific attraction to chemotypes, we conducted binomial tests to test if aphids preferred one chemotype over another for each pairwise combination (see supplement 1.2). We tested whether the number of choices made for one chemotype in a specific combination was significantly different from what would be expected in a random choice. Binomial tests were also used to rule out spatial (left/right side of the petri dish) effects on aphid preference. We ruled out observer effects on the choices, as well as on the proportion of choice/no-choice using Chi-square tests. Clogit models were used to investigate the attractiveness of specific tansy chemotypes and daughters for each aphid species, across all combined pairwise tests with all replicates (see Table 2 & 3), using the survival package (Therneau 2021). We used the z-values obtained from the model for each tansy daughter as a proxy for attractiveness, with daughters with more positive z-values being more and negative z-values being less attractive to aphids.

To test our second hypothesis, whether chemodiversity negatively correlated with attractiveness, we calculated plant chemodiversity metrics (compound richness, Shannon diversity, evenness, approximate total concentration) from the absolute terpenoid profiles, using the vegan package (Oksanen et al. 2020) and tested them against aphid preference. We tested for relationships between the approximate total concentrations of individual compounds in a plant and attractiveness of the plant separately for each of the two aphid species. We used Spearman correlations, as changes in chemical compounds within individual profiles are potentially related to one another as they might occur on the same synthetic pathways, but between individuals these relationships might not be linear. We analysed correlations of terpenoids and aphid preference at the tansy daughter level (n = 18, note: we did not have data for the three daughters from Aue41 regarding aphid preference due to propagation issues). We used unadjusted correlation plots and Holm-adjusted plots for multiple correlations, using the RcmdrMisc package (Fox 2022) and present both for visualization purposes. For verification purposes, we ran a multiple regression model testing the effect of all compounds on attractiveness. As the models were limited by available degrees of freedom, we removed all compounds that were measured in less than two-thirds of all samples. Using step() in combination with variance inflation factors to address multicollinearity, we reduced the model to the minimum adequate version, which only included individual compounds and proved out to be non-significant.

To test our third hypothesis, whether there were growth-related trade-offs with chemodiversity, we tested the height, number of stems, number of leaves, leaflets density, specific leaf area and chlorophyll against terpenoid richness, terpenoid Shannon diversity, terpenoid Shannon evenness and approximate total terpenoid concentration, by using one-way ANOVA with chemotype as a fixed factor (six chemotypes, with n = 3 daughters).

## Results

### Chemical profile of chemotypes

Selected chemotypes differed based on their terpenoid blend composition (Fig. 2a), as well as terpenoid diversity, evenness, richness and approximate total concentration (Fig. 2b-e). Specifically, Aue41 and Schienen3 were both dominated by β-thujone. Schie5 had both, α- and β-thujone as prevalent compounds. Most of the terpenoid blend in Saal2 was made up by β-trans-chrysanthenyl acetate. In the chemotype Spar11B, sabinene and α-pinene were made up of 39% and 20% of the terpenoid concentration, but no terpenoid was clearly dominating. The chemical profile of Cam7 was very mixed with no compound clearly dominating. Cam7 and Spar11B had significantly higher terpenoid Shannon diversity and terpenoid evenness than other chemotypes (Fig. 2b-c). The chemotype did not significantly affect chemotype richness (Fig. 2d). Spar11B and Aue41 had significantly lower approximate total terpenoid concentrations compared to the other chemotypes (Fig. 2e).

**Figure 2:**
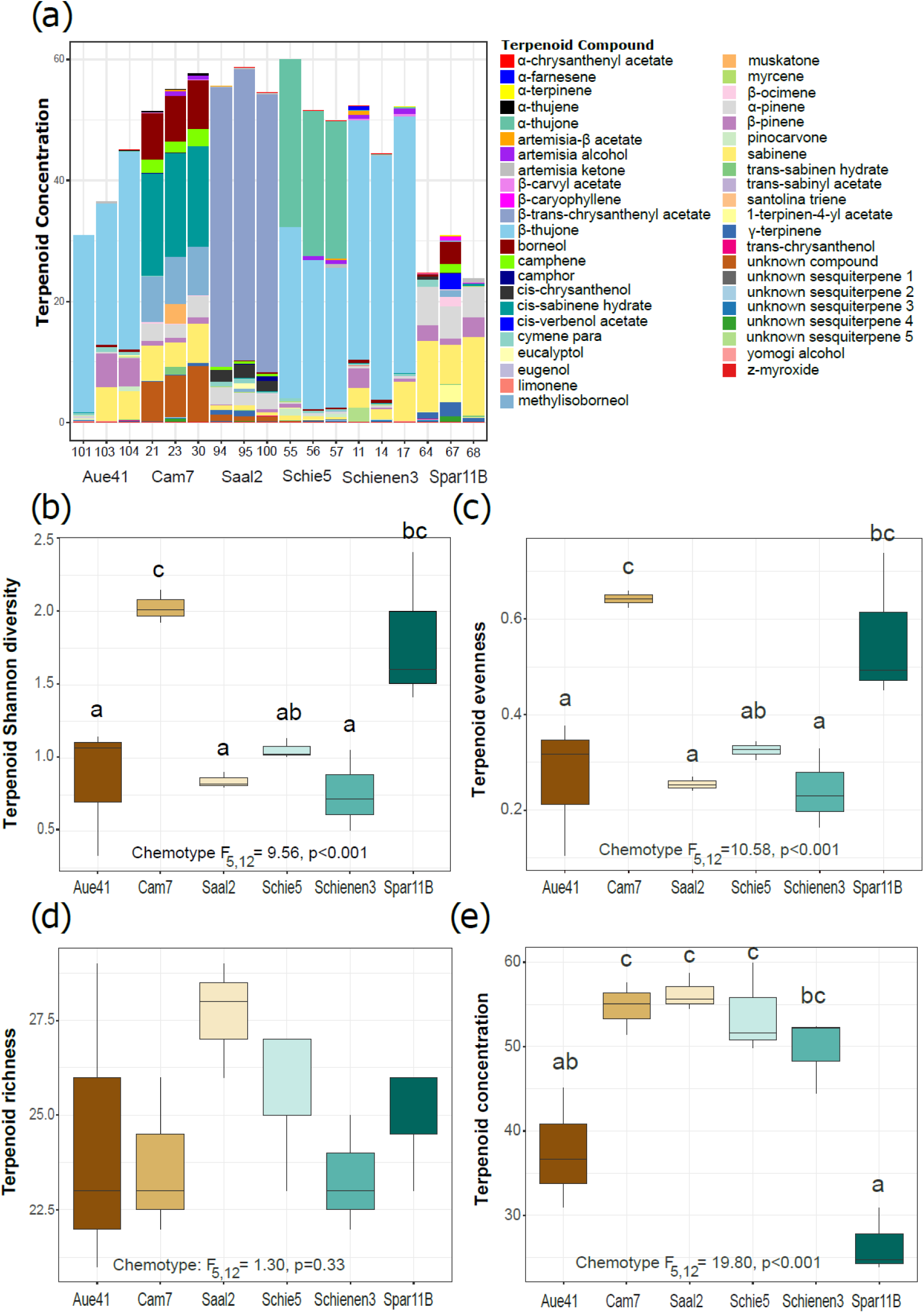
a) Stacked bar chart chemical profiles for the selected six maternal chemotypes with three daughter replicates. Box plots show the interquartile ranges of b) terpenoid Shannon diversity, c) terpenoid evenness, d) terpenoid richness, and e) approximate total terpenoid concentration for each chemotype. The lower hinge corresponds to the first quartile (25^th^ percentile) and the upper hinge depicts the third quartile (75^th^ percentile). Whiskers extend to the 5% and 95% percentiles; solid lines represent the medians. Chemotype effect is indicated in panels, letters above bars indicate significant differences (p<0.05) between chemotypes based on post-hoc Tukey tests.

### Choice experiments

After two hours, seven *M. tanacetaria* and five *U. tanaceti* individuals were found dead and were excluded from the analyses. 178 *M. tanacetaria* and 69 *U. tanaceti* had chosen a specific chemotype pinna. *M. tanacetaria* aphids tended to marginally prefer leaflets from Schie5 over leaflets from Spar11B and Schienen3 (Table 1, Fig. 3a). After five hours of observation, *M. tanacetaria* showed a significant preference for Schie5 over Spar11B and tended to prefer Saal2 over Cam7 and Spar11B (Table 1, Fig. 3b). After two hours, *U. tanaceti* aphids significantly preferred Schie5 over Spar11B (Table 1, Fig. 3c), but after five hours no more significant preferences were observed (Table 1, Fig. 3d). Considering all aphid choices made across all pairwise comparisons in a clogit model, *M. tanacetaria* aphids showed a significant attraction to Saal2 and a marginally significant attraction to Schie5 chemotypes (Table 2, Fig. 4a), while *U. tanaceti* aphids did not show a clear preference for any chemotype (Table 2, Fig. 4b,). However, we found that aphids also exhibited preferences at the individual plant daughter level, with *M. tanacetaria* showing a significant preference for Saal2-95 and Saal2-100, and Schie5-55 and Schie5-56, whereas *U. tanaceti* did not show significant preferences to any tansy daughters (Table 3).

**Figure 3:**
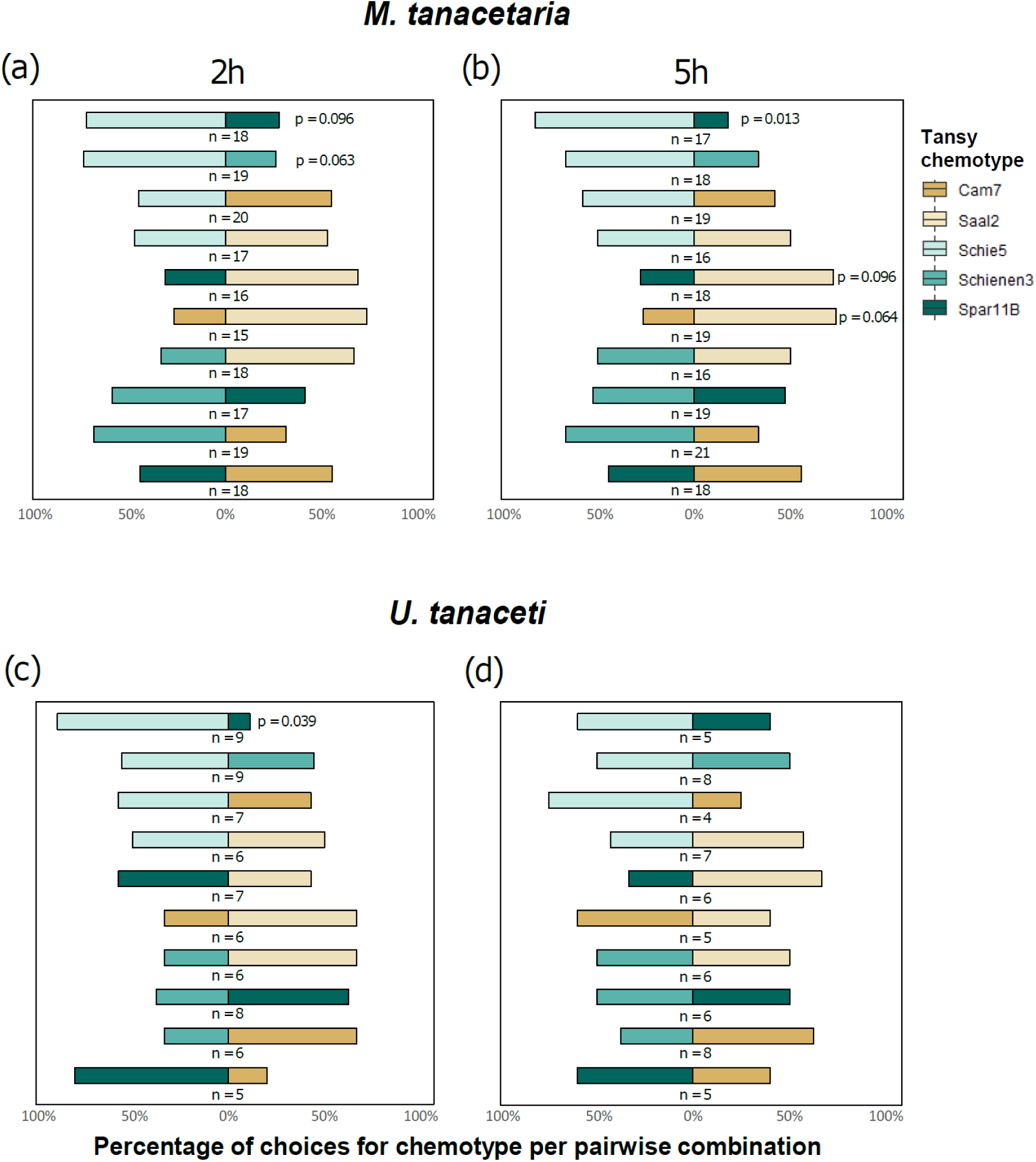
Mean percentage of choices by a) *Macrosiphoniella tanacetaria*, and b) *Uroleucon tanaceti* after two hours for all possible pairwise combinations of chemotypes, with error bars indicating standard errors. Effective sample size is indicated for each pairwise comparison, and no-choice replicates were excluded. Note that chemotype Aue41 was not included in pairwise comparisons due to propagation difficulties. P-values next to bars indicate (marginally) significant preferences (for test statistics see Table 1).

**Figure 4:**
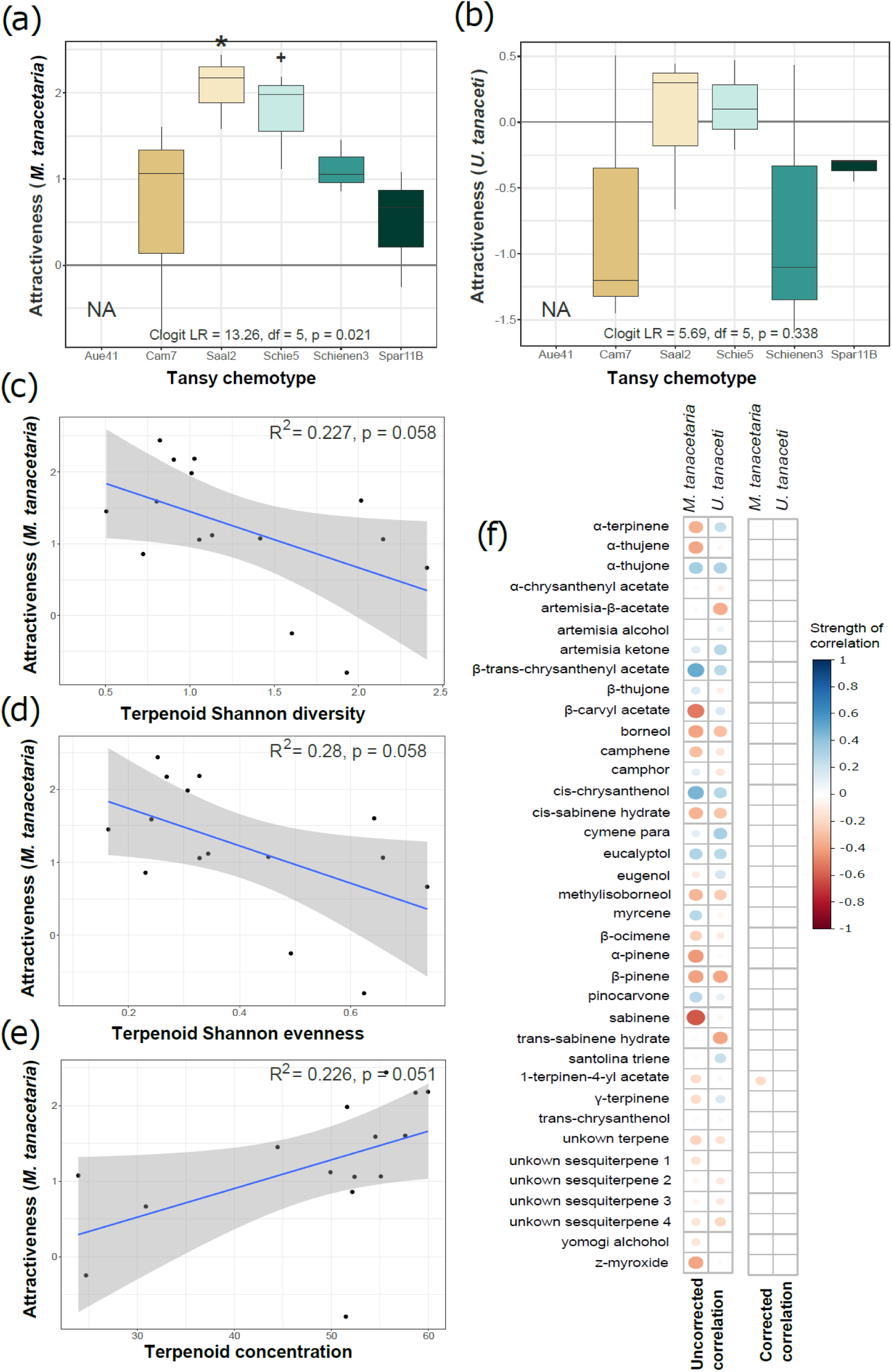
Relationships between tansy attractiveness and chemodiversity for both aphid species. Attractiveness (reflected by the z-value) was computed by summarizing all decisions made towards a specific chemotype over all combinations and using a clogit model to test whether a certain chemotype was chosen more often than expected compared to a random choice. a) mean attractiveness per chemotype for *M. tanacetaria.* b) Mean attractiveness per chemotype for *U. tanaceti*. Symbols indicate significance levels (* p < 0.05; + 0.05 < p < 0.10). For detailed model output for a) and b) see Table 2. c) Relationship between tansy terpenoid Shannon diversity and attractiveness to *M. tanacetaria*. d) Relationship between tansy terpenoid evenness and attractiveness to *M. tanacetaria*. e) Relationship between tansy terpenoid abundance and attractiveness to *M. tanacetaria*; in c) to e) dots represent the mean z-value (attractiveness) for each of the 15 daughters (5 Chemotypes * 3 daughters). f) Correlation plots showing relationship between individual terpenoid compounds and attractiveness to both aphid species. The size of the dot depicts the strength of the correlation. Blue colours indicate positive and red colours indicate negative correlations. The left correlation plot shows all unadjusted correlations with a significant p-value. The right correlation plot shows all correlations with a significant p-value after using Holm-adjusted for multiple correlations.

**Table 1:** Influence of tansy chemotypes on their attractiveness to *M. tanacetaria* and *U. tanaceti*, when given pairwise choices. A binomial test was used to obtain credible intervals and p-values. Significant values are highlighted in bold, marginally significant values in italics.

| Time | Chemotype 1 | Chemotype 2 | *M. tanacetaria* |  |  | *U. tanacetii* |  |  |
| --- | --- | --- | --- | --- | --- | --- | --- | --- |
|  |  |  | n | 95% CI | p-value | n | 95% CI | p-value |
| 2 h | Schie5 | Spar11B | 18 | *0.465, 0.903* | *0.096* | 9 | **0.518, 0.997** | **0.039** |
| 2 h | Schie5 | Schienen3 | 19 | *0.488, 0.909* | *0.064* | 9 | 0.212, 0.863 | 1 |
| 2 h | Schie5 | Cam7 | 20 | 0.231, 0.685 | 0.824 | 7 | 0.184, 0.901 | 1 |
| 2 h | Schie5 | Saal2 | 17 | 0.230, 0.722 | 1 | 6 | 0.118, 0.882 | 1 |
| 2 h | Spar11B | Saal2 | 16 | 0.110, 0.587 | 0.210 | 7 | 0.184, 0.901 | 1 |
| 2 h | Cam7 | Saal2 | 15 | 0.078, 0.51 | 0.119 | 6 | 0.043, 0.778 | 0.688 |
| 2 h | Schienen3 | Saal2 | 18 | 0.133, 0.590 | 0.238 | 6 | 0.043, 0.778 | 0.688 |
| 2 h | Schienen3 | Spar11B | 17 | 0.330, 0.816 | 0.629 | 8 | 0.085, 0.755 | 0.727 |
| 2 h | Schienen3 | Cam7 | 19 | 0.435, 0.874 | 0.167 | 6 | 0.043, 0.778 | 0.688 |
| 2 h | Spar11B | Cam7 | 18 | 0.215, 0.692 | 0.815 | 5 | 0.284, 0.995 | 0.375 |
| 5 h | Schie5 | Spar11B | 17 | **0.566, 0.962** | **0.013** | 5 | 0.147, 0.947 | 1 |
| 5 h | Schie5 | Schienen3 | 18 | 0.410, 0.867 | 0.238 | 8 | 0.157, 0.843 | 1 |
| 5 h | Schie5 | Cam7 | 19 | 0.335, 0.798 | 0.648 | 4 | 0.194, 0.994 | 0.625 |
| 5 h | Schie5 | Saal2 | 16 | 0.247, 0.754 | 1 | 7 | 0.099, 0.816 | 1 |
| 5 h | Spar11B | Saal2 | *18* | *0.097, 0.535* | *0.096* | 6 | 0.043, 0.778 | 0.688 |
| 5 h | Cam7 | Saal2 | *19* | *0.092, 0.512* | *0.064* | 5 | 0.147, 0.947 | 1 |
| 5 h | Schienen3 | Saal2 | 16 | 0.247, 0.754 | 1 | 6 | 0.118, 0.882 | 1 |
| 5 h | Schienen3 | Spar11B | 19 | 0.289, 0.756 | 1 | 6 | 0.118, 0.882 | 1 |
| 5 h | Schienen3 | Cam7 | 21 | 0.430, 0.854 | 0.189 | 8 | 0.085, 0.755 | 0.727 |
| 5 h | Spar11B | Cam7 | 18 | 0.215, 0.692 | 0.815 | 5 | 0.147, 0.947 | 1 |

**Table 2:** Influence of tansy chemotypes on their attractiveness to *M. tanacetaria* and *U. tanaceti* based on all pairwise combinations of the choice assays after two hours of observation. A c-logit model, including effects of chemotype (five chemotypes) on aphid choices after two hours, was used to obtain z- and p-values (for visualization see figure 4). Significant values are highlighted in bold, marginally significant values in italics.

| Chemotype | *M. tanacetaria* |  | *U. tanaceti* |  |
| --- | --- | --- | --- | --- |
|  | z-value | p-value | z-value | p-value |
| Cam7 | -0.669 | 0.503 | -1.226 | 0.220 |
| Saal2 | **2.331** | **0.020** | 0.783 | 0.434 |
| Schie5 | *1.730* | *0.084* | 1.206 | 0.228 |
| Schienen3 | 0.340 | 0.734 | -0.448 | 0.654 |
| Spar11B | -0.752 | 0.452 | 0.404 | 0.686 |

**Table 3:**
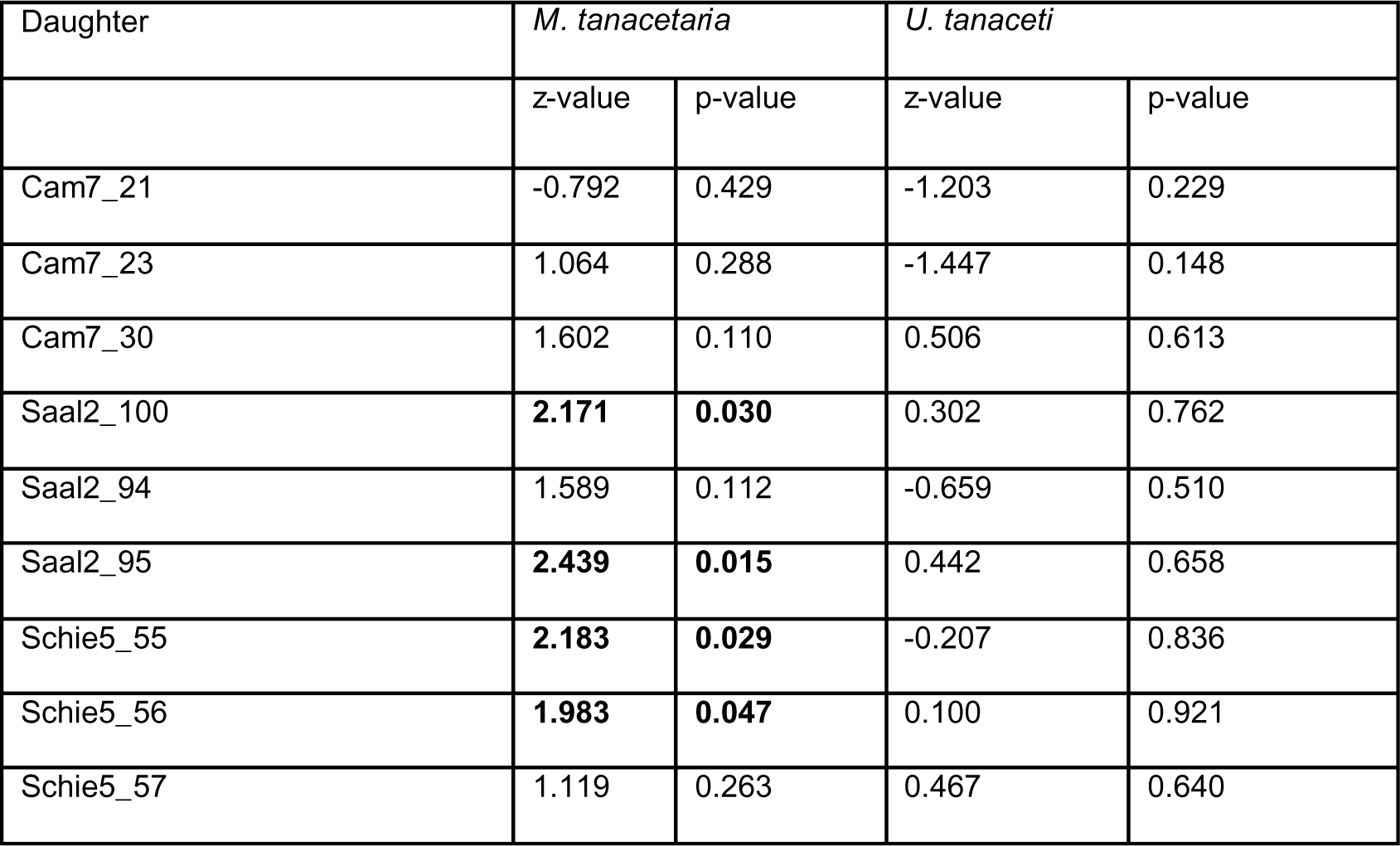
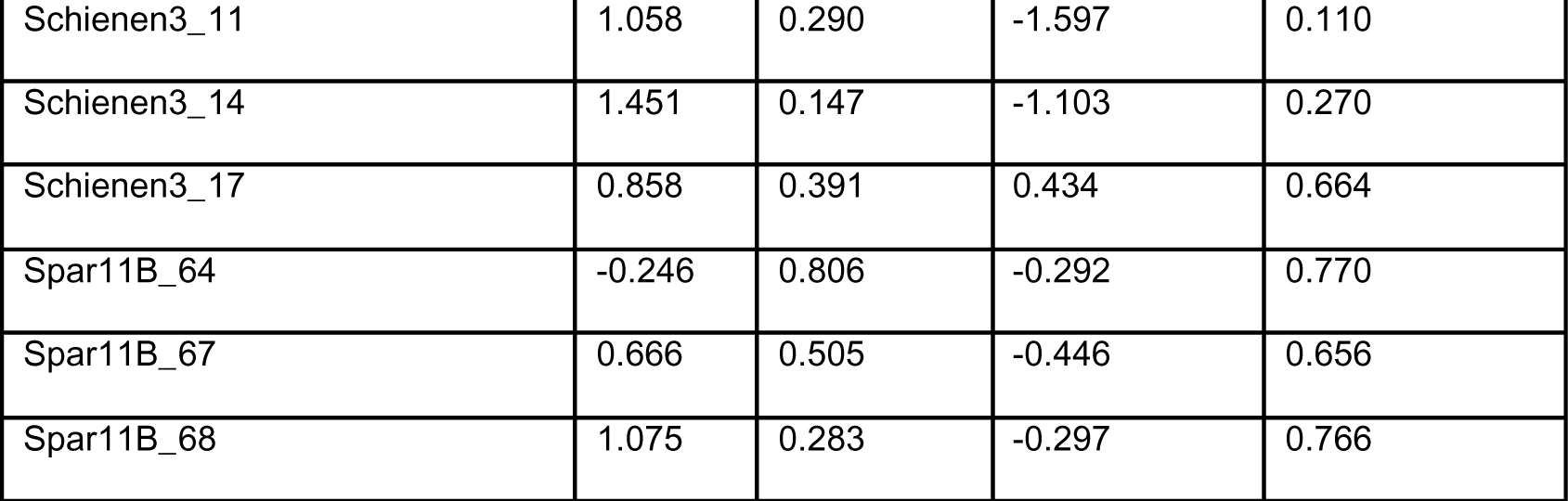
Influence of tansy daughters on their attractiveness to *M. tanacetaria* and *U. tanaceti* based on all pairwise combinations of the choice assays after two hours of observation. A c-logit model, including effects of daughter (five chemotypes x three daughters) on aphid choices after two hours, was used to obtain z- and p-values. Significant values are highlighted in bold, marginally significant values in italics.

| Daughter | *M. tanacetaria* |  | *U. tanaceti* |  |
| --- | --- | --- | --- | --- |
|  | z-value | p-value | z-value | p-value |
| Cam7_21 | -0.792 | 0.429 | -1.203 | 0.229 |
| Cam7_23 | 1.064 | 0.288 | -1.447 | 0.148 |
| Cam7_30 | 1.602 | 0.110 | 0.506 | 0.613 |
| Saal2_100 | **2.171** | **0.030** | 0.302 | 0.762 |
| Saal2_94 | 1.589 | 0.112 | -0.659 | 0.510 |
| Saal2_95 | **2.439** | **0.015** | 0.442 | 0.658 |
| Schie5_55 | **2.183** | **0.029** | -0.207 | 0.836 |
| Schie5_56 | **1.983** | **0.047** | 0.100 | 0.921 |
| Schie5_57 | 1.119 | 0.263 | 0.467 | 0.640 |
| Schienen3_11 | 1.058 | 0.290 | -1.597 | 0.110 |
| Schienen3_14 | 1.451 | 0.147 | -1.103 | 0.270 |
| Schienen3_17 | 0.858 | 0.391 | 0.434 | 0.664 |
| Spar11B_64 | -0.246 | 0.806 | -0.292 | 0.770 |
| Spar11B_67 | 0.666 | 0.505 | -0.446 | 0.656 |
| Spar11B_68 | 1.075 | 0.283 | -0.297 | 0.766 |

For *M. tanacetaria*, we observed a trend of decreased attraction to plants with higher diversity (Fig. 4c) and more evenly distributed terpenoid blends (Fig. 4d), but a trend of increased attraction to plants with a higher approximate total terpenoid concentration (Fig. 4e). For *U. tanaceti*, we observed no relationships between attractiveness and chemodiversity. Unadjusted correlation plots revealed that various relationships existed between individual compounds and attractiveness of aphid species, but when Holm-adjusted for multiple correlations were applied, this reduced the relationships between attractiveness and individual terpenoids to only a weak negative relationship between 1-terpinen-4-yl acetate and tansy attractiveness to *M. tanacetaria* (Fig. 4f). This was further verified with simplified multiple regression models corrected for collinearity, which reduced the models to single compound factors, which did not significantly affected attractiveness.

### Correlation between chemical diversity and morphological plant traits

Significant positive relationships were observed between the number of stems per tansy plant and terpenoid Shannon diversity (Table 4, Fig. 5a), tansy height and terpenoid Shannon diversity (Fig. 5b), and tansy height and approximate total terpenoid concentration (Fig. 5c).

**Figure 5:**
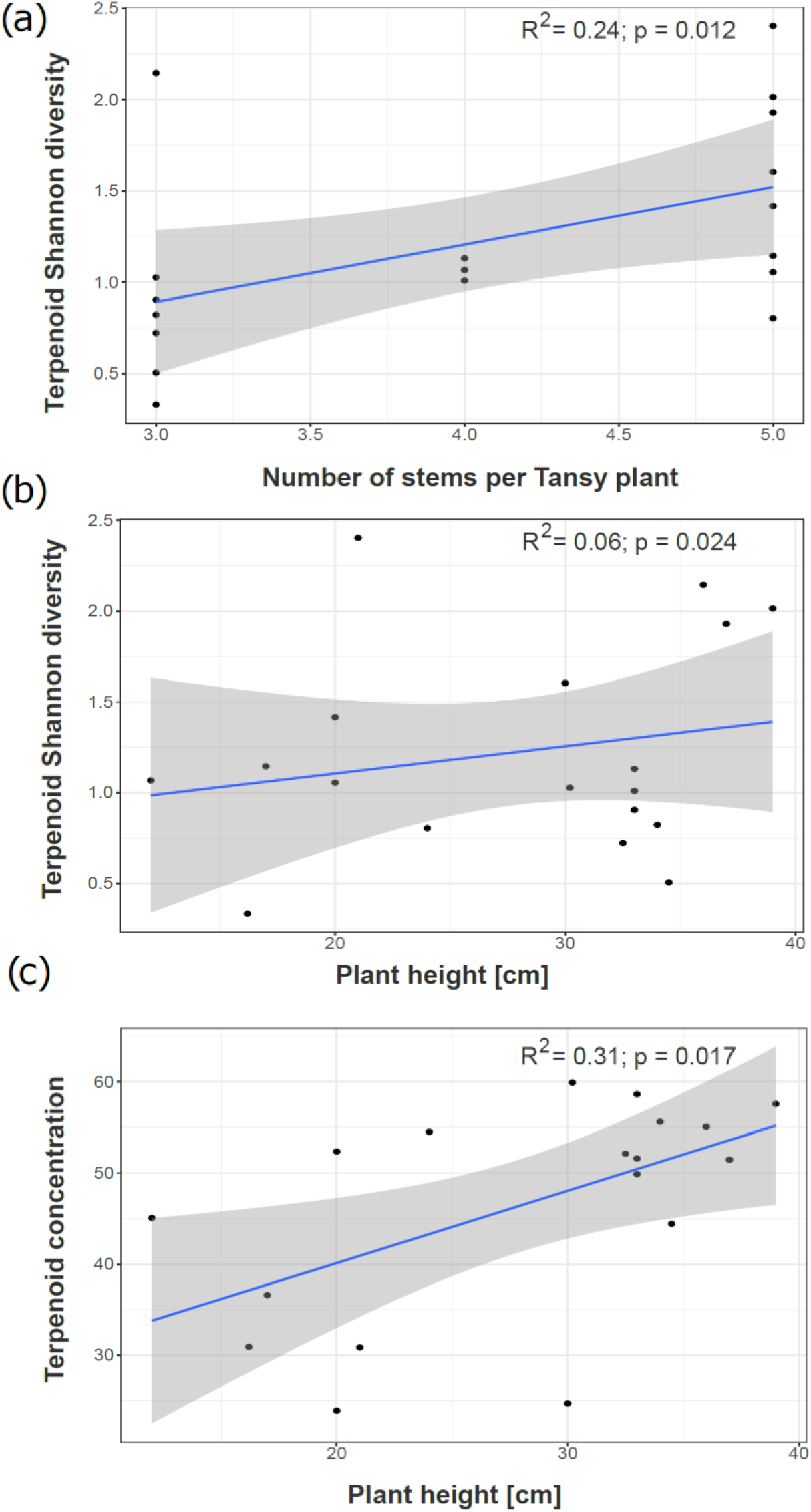
Relationships between chemical and morphological parameters. Each dot represents one of the 18 daughters (6 chemotypes * 3 daughters). a) Relationship between the number of stems per tansy plant and terpenoid Shannon diversity. b) Relationship between tansy height and terpenoid Shannon diversity. c) Relationship between tansy height and approximate total terpenoid concentration.

**Table 4:** Model output of multiple linear regression models testing the effects of plant morphological characteristics on plant attractiveness for *M. tanacetaria* and *U. tanaceti*, and on plant chemodiversity characteristics. Models were checked for collinearity by variance inflation factors and simplified via stepwise model selection. Dashes indicate variables were removed from the model.

| Explanatory variable | Attractiveness *M. tanacetaria* | Attractiveness *U. tanaceti* | Terpenoid richness | Terpenoid Shannon diversity | Terpenoid Shannon evenness | Terpenoid concentration |
| --- | --- | --- | --- | --- | --- | --- |
|  | F (p-value) | F (p-value) | F (p-value) | F (p-value) | F (p-value) | F (p-value) |
| Height | - | - | 0.20 (0.665) | **6.03 (0.028)** | **6.44 (0.024)** | **7.05 (0.017)** |
| # Stems | **10.20 (0.010)** | - | 0.11 (0.741) | **8.33 (0.012)** | **9.12 (0.009)** | - |
| # Leaves | **5.02 (0.049)** | - | 0.07 (0.801) | - | - | - |
| Leaflets density | 3.14 (0.107) | 1.85 (0.199) | - | - | - | - |
| SLA | - | - | - | - | - | - |
| Chlorophyll | **6.94 (0.0250)** | 2.42 (0.146) | - | 2.55 (0.133) | 2.54 (0.133) | - |

## Discussion

In this study, we found that tansy plants clustered into six distinct chemotypes, based on their terpenoid profile. Chemotypes furthermore differed in their terpenoid Shannon diversity, terpenoid evenness and terpenoid concentration. We used five distinct maternal chemotype lines to test how leaf chemical profiles affected aphid preference in pairwise choice assays. We found that two aphid species specialized on tansy showed species-specific preferences in some of the pairwise choices. Across all pairwise combinations *Macrosiphoniella tanacetaria* showed stronger preferences and patterns of attraction than *Uroleucon tanaceti*. In line with our expectations, we observed a trend of higher attractiveness of plants with a higher approximate total terpenoid concentration and lower diversity. These patterns were observed for *M. tanacetaria*, but not for *U. tanaceti*. Interestingly, attraction of aphids could not be clearly linked to individual terpenoids.

Supporting our first hypothesis, we show that aphids preferred different chemotypes in pairwise choice assays. *M. tanacetaria* preferred leaves from plants with α-thujone/β-thujone (Schie5) and β-trans-chrysanthenyl acetate (Saal2) chemotype, with preference becoming more pronounced after five hours. Previous studies found contradicting results, with higher numbers of *M. tanacetaria* on plants with β-thujone compared to trans-carvyl acetate in a climate chamber experiment (Jakobs & Müller 2018), but a higher abundance of *M. tanacetaria* on plants with camphor instead of β-thujone as dominant compound in a field study (Benedek et al. 2019a). Similarly, *U. tanaceti* was significantly attracted towards Schie5 (α-thujone/β-thujone), with preference becoming less pronounced over time. This is interesting, as previously negative effects of β-thujone on *U. tanaceti* numbers have been found in the field (Benedek et al. 2019b), although no clear preference was observed towards chemotypes for this species in another common garden study (Kleine & Müller 2011). Under field conditions different factors (such as insect preference, bottom-up and top-down processes) affect aphid communities simultaneously, and this may explain some of the discrepancies between field observations and choice assays, as in ‘snapshot’ field observations, these factors can be hard to disentangle. Therefore, studies under laboratory conditions are necessary to understand the influence of these single factors on aphid behaviour and performance. Our results indicate that aphid preference for plants might not always reflect how they perform on plants.

In our experiment, 70 - 91 % of *M. tanacetaria* individuals made a choice after five hours for each combination, compared to only 30 - 61% of *U. tanaceti* individuals. These species-specific differences between the aphids can possibly be explained by their life history and the difference in their preferred niche (Jakobs, Schweiger & Müller 2019), and aligns with our observations. We observed in various field and greenhouse experiments that these aphid species showed different behaviours. For instance, *M. tanacetaria* tends to be more mobile than *U. tanaceti*, and readily searches for new host plants when its current host deteriorates. *Uroleucon tanaceti* typically remains on leaves until yellowing, to then move upward to a new leaf, often directly above the one they were on. We found more deaths and fewer choices made at later time points (i.e., after 24 h), which could be because individual leaflets do dry out after some time. However, as individual choices were already apparent and similar after two and five hours, we believe that deteriorating quality did not affect choices made after two hours.

Across all experimental choice combinations, we found significant effects of chemotype on attractiveness of those chemotypes for *M. tanacetaria*, but not for *U. tanaceti*. This is in line with previous research from Kleine & Müller (2011), who found that *M. tanacetaria* did, but *U. tanaceti* did not exhibit distinct preferences towards specific chemotypes. To understand what the drivers of the attractiveness of a plant to *M. tanacetaria* may be, we investigated relationships between tansy individual chemical properties and their level of attractiveness. We observed that diversity of the terpenoid blend negatively correlated with attractiveness. This is consistent with the hypothesis on the evolution of chemodiversity in plants, and particularly the evolution of a breadth of specialized compounds to repel antagonists (Wetzel & Whitehead 2020). Furthermore, we found that terpenoid evenness marginally negatively correlated with attractiveness, indicating that blends that were more evenly distributed in terpenoid composition were less attractive, than those dominated by one or several compounds. A plausible explanation might be that having some highly dominant compounds could serve as strong cues for plant recognition (Kleine & Müller 2011). Subsequent aphid performance on tansy likely is affected more strongly by phloem composition (Jakobs & Müller 2019). We also found a positive effect of approximate total terpenoid concentration on attractiveness to *M. tanacetaria*. This might be an indication that although chemodiversity can have a deterrent effect (Whitehead et al. 2021), the approximate concentration (as a proxy for potential emission) of terpenoids can be an important cue for host finding. A role of specialized metabolite concentration in attraction and repellence has been found in other studies (reviewed in Macel 2011). For instance, contrasting effects have been observed in the specialized aphid *Aphis jacobaea* on *Jacobaea vulgaris*, where plants high in pyrrolizidine alkaloids hosted fewer aphids than plants with low concentrations, although this could not be related to preference alone (Vrieling et al. 1991). Our results align with those from a recent study that shows that chemodiversity is an important driver of dietary specialization in insects (Leong et al. 2022). The relationships between *M. tanacetaria* and approximate total terpenoid concentration, terpenoid diversity and evenness were marginally significant, which was likely a result of the limiting number of chemotype lines (15) included in our current study. Our study was limited by the poor propagation success of one of the chemotype lines with replicates. It is likely that including a larger number of diverse chemotypes would further strengthen these findings, and future studies on a range of terpenoid diversity profiles would be meaningful.

Our analyses of the role of individual compounds in driving attractiveness indicated that the effect of individual compounds as drivers of tansy attractiveness are minimal. After corrections for multiple correlations, we observed a weak negative effect of 1-terpinen-4-yl acetate on tansy attractiveness to *M. tanacetaria*. Several studies have shown relationships between chemotypes, or dominant compounds and aphid abundance in the field. For instance, β-thujone has been related to decreased colony distribution and colony numbers in *U. tanaceti* and abundances in *M. tanacetaria* in field studies (Bálint et al. 2016, Benedek et al. 2019b). Other studies have found an increased density of *Metopeurum fuscoviride* on chemotypes with high concentrations of borneol or camphor (Bálint et al. 2016, Senft et al. 2019), while plants with high amounts of α-thujone, (E)-dihydrocarvone, α-copaene and β-cubebene were colonized earlier (Clancy et al. 2016). However, importantly, all these studies investigated presence and abundance of aphids in the field, and did not take into account aphid preference. Hence these studies are limited in the information they can provide about aphid choice. Although we find only limited evidence that individual compounds drive aphid attractiveness, it cannot be ruled out that these compounds play a role in feeding deterrence when aphids are on the plant, or attraction of herbivore antagonists. Importantly, our study included terpenoids only, but other volatile compounds (e.g., flavonoids) that were not detected here, may affect the preference behavior of the aphids. This calls for future studies disentangling the role of broader chemodiversity in driving aphid performance predator attraction.

It is important to disentangle the relation between chemical and growth traits, as both traits have been shown to influence plant-insect interactions, including preference. While growth-defence trade-offs are a fundamental principle in plant ecology (Herms & Mattson 1992; Karasov et al. 2017), we did not observe such a trade-off in our model system. Instead, we found a synergistic effect between variables associated with plant growth, and approximate total terpenoid concentration and diversity in our plants. It is commonly assumed that plants have to partition their resources between growth and defence traits, leading to either smaller and better defended plants, or *vice versa* (Coley at al. 1985, Herms & Mattson 1992, He et al. 2022). Terpenoids are more expensive to build than many other metabolites, as they require a wide array of different enzymes, hence posing substantial production and storage costs (Gershenzon 1994). However, contrary to these hypotheses, we found that larger and bushier plants have a higher approximate total concentration and Shannon diversity of terpenoid compounds. Similarly, Barton in 2007 found a positive relationship of growth and defense in two *Plantago* species. One plausible explanation for our findings may be that larger plants photosynthesize more, and have a larger energy budget, which can be used for elevated and diversified local terpenoid synthesis. Furthermore, it is vital for plants to grow *and* defend to optimize their fitness within a dynamic environment (Huot et al. 2014), and trade-offs might also change with different life ontogeny stages (Boege & Marquis 2005). Recent work has shown strong effects of maternal “chemo-genotypes” on plant chemical composition in tansy (Dussarrat et al. 2023). Given that our chemotypes were confounded by their maternal origin (chemo-genotypes), this may also affect chemical and morphological traits, and future work should investigate whether chemo-genotypes also influences aphid attraction and performance.

In this study, we used unwinged aphid morphs. Although in early colonization in the field, winged morphs are generally more likely to choose host plants than unwinged morphs (Mehrpavar et al. 2014), dispersal to other plants is commonly observed for unwinged aphids in this system, both in experimental colonies and the field (the authors, pers. obs.). Although performing choice assays with winged early colonizers would be an important next step, such early wing morphs are difficult to obtain experimentally. In a choice assay study, Mehrparvar and colleagues (2014) found that winged dispersal morphs of *M. tanacetaria* displayed preferences for plants with specific herbivore infestation history, while unwinged aphids did not show any such host plant preferences. Our results indicate that preferences and choices in unwinged morphs do occur at the chemotype level, which likely represents a stronger chemical contrast for aphids than the herbivory history studied in previous work.

Even though aphids preferred specific chemotypes, these might not necessarily be the most suitable chemotypes for colony development. There has been a whole debate regarding the “mother knows best” principle which is connected to the preference-performance hypothesis, which posits that females insects prefer to oviposit on host plants on which their offspring performs best (Martínez et al 2013, Gripenberg et al 2010). In the beetle *Gonioctena linnaeana* for example, it has been found that mothers preferred host plants on which the survival rate of their larvae was significantly higher (Wennström et al. 2010), while females of the root-feeding vine weevil *Otiorhynchus sulcatus* preferred to oviposit on plants with small root systems, which decreased offspring performance (Clark et al. 2011). Whether preference overlaps with performance differs between species (Valladares & Lawton 1991, García-Robledo & Horvitz 2012). For *U. tanaceti* the “mother knows best” principle seems to apply, as they chose the chemotype and plant parts in a choice-study, on which their offspring performed best, while in *M. tanacetaria* this was not the case (Jakobs & Müller 2018).

## Conclusion

In this study, we found that plant chemodiversity is a driver for aphid host plant preference, which is in line with other studies (Wolf et al. 2012), strengthening our understanding of the role of chemodiversity in plants. We found that different chemotypes have distinct attraction patterns, and that these can be explained in part by diversity metrics of the terpenoid blend, as well as the approximate total concentration. Insects use chemical cues, including host plant metabolites, to inform their decisions. It is eminent that we develop a better understanding of how intraspecific plant chemodiversity shapes the various aspects of the (herbivorous) insect life cycle, including development, survival, defense, and overall fitness.

## Acknowledgements

We thank the staff at TUM PTC Dürnast, particularly Sabine Zuber and Petra Scheuerer, for providing excellent care during plant propagation and preparation phase. We want to thank the DFG for funding. This study was funded by the German Research Foundation (DFG), project 415496540 to WWW (WE3081/40-1) and to CM (MU1829/28-1), as part of the Research Unit (RU) FOR 3000. Furthermore, it was supported by DFG project 245400135 to WWW (WE3081/25-2).

## Contribution of authors

ANH, LOP, WWW and RH conceived and designed the experiment. LOP and RH propagated plant material. EE and CM chemotyped plant material. RH and LOP performed morphological trait measurements. ANH and LOP performed the choice assays. ANH, LOP and RH analysed the data. ANH and RH wrote the paper, with substantial input from LOP. All authors read and approved the manuscript before submission.

