## Supplementary information for "Chemodiversity affects preference for *Tanacetum vulgare* chemotypes in two aphid species"

1. R-code of statistical models used

1.1 Clustering of plants into chemotypes

D <- vegdist(data, method = “Euclidean”)

Hclust(D, method = “ward.D2”)

1.2 Binomial tests for aphid preference towards in pairwise choice assay

Binom.test(chosen, not chosen, p=0.5, alternative = “two-sided”, confidence level = 0.95)

1.3 Attractiveness towards chemotypes and daughters overall

clogit(Chosen(yes, no) ~ Daughter + Side (left, right) + strata(ID of aphid), data = decisions of *M. tanacetaria*/*U. tanaceti* after 2 hours)

1.4 Plant diversity metrics

Shannon diversity: H <- diversity(data)

Compound richness: S <- specnumber(data)

Evenness: J <- H/log(S)

Anova(lm(preference~shannon + evenness + richness))

1.5 Correlation of single compounds and aphid preference

Data_cor <- Cor(data terpenoids, data z-value preference for each species, use = “pairwise complete observations”)

Corrected p-value: using RcmdrMisc package (Fox 2022)

rcorr.adjust(Data_cor, type =”spearman”, use=”pairwise complete observations”)

(Holm correction by default)

1.6 Multiple regression for verification

Model <- lm(attractiveness ~all terpenoid compounds found in 10 out of 15 daughters (>66%))

vif(Model) #calculates the variation inflation factors. Terms with >1.5 exhibit multicollinearity and therefore are removed from the model

step(Model) #chooses stepwise the best model

1.7 One-way ANOVA for correlation of plant growth and chemodiversity

Model <- (lm(shannon diversity ~height + number stems + number leaves +

pinnae density + specific leaf area + chlorophyll))

Vif(model)

Step(model)

Anova(lm(shannon diversity ~ height + number stems + chlorophyll))

Table S1: The number of replicates for the combinations of daughters used in the experiment, both for *M. tanacataria* and *U. tanaceti*.

| Replicates for *M. tanacetaria* | Replicates for *U. tanaceti* | Daughter 1 | Daughter 2 |
|---|---|---|---|
| 3 | 3 | Schienen3_11 | Cam7_21 |
| 2 | 2 | Schienen3_11 | Cam7_23 |
| 2 | 0 | Schienen3_11 | Cam7_30 |
| 3 | 3 | Schienen3_11 | Schie5_55 |
| 2 | 2 | Schienen3_11 | Schie5_56 |
| 2 | 0 | Schienen3_11 | Schie5_57 |
| 3 | 3 | Schienen3_11 | Spar11B_64 |
| 2 | 2 | Schienen3_11 | Spar11B_67 |
| 2 | 0 | Schienen3_11 | Spar11B_68 |
| 3 | 3 | Schienen3_11 | Saal2_94 |
| 2 | 2 | Schienen3_11 | Saal2_95 |
| 2 | 0 | Schienen3_11 | Saal2_100 |
| 2 | 2 | Schienen3_14 | Cam7_21 |
| 3 | 1 | Schienen3_14 | Cam7_23 |
| 3 | 1 | Schienen3_14 | Cam7_30 |
| 2 | 2 | Schienen3_14 | Schie5_55 |
| 3 | 1 | Schienen3_14 | Schie5_56 |
| 3 | 1 | Schienen3_14 | Schie5_57 |
| 2 | 2 | Schienen3_14 | Spar11B_64 |
| 3 | 1 | Schienen3_14 | Spar11B_67 |
| 3 | 1 | Schienen3_14 | Spar11B_68 |
| 2 | 2 | Schienen3_14 | Saal2_94 |
| 3 | 1 | Schienen3_14 | Saal2_95 |
| 3 | 1 | Schienen3_14 | Saal2_100 |
| 2 | 0 | Schienen3_17 | Cam7_21 |
| 3 | 1 | Schienen3_17 | Cam7_23 |
| 3 | 3 | Schienen3_17 | Cam7_30 |
| 2 | 0 | Schienen3_17 | Schie5_55 |
| 3 | 1 | Schienen3_17 | Schie5_56 |
| 3 | 3 | Schienen3_17 | Schie5_57 |
| 2 | 0 | Schienen3_17 | Spar11B_64 |
| 3 | 1 | Schienen3_17 | Spar11B_67 |
| 3 | 3 | Schienen3_17 | Spar11B_68 |
| 3 | 1 | Schienen3_17 | Saal2_94 |
| 3 | 1 | Schienen3_17 | Saal2_95 |
| 2 | 2 | Schienen3_17 | Saal2_100 |

| | | | |
|---|---|---|---|
| 5 | 3 | Cam7_21 | Schie5_55 |
| 2 | 2 | Cam7_21 | Schie5_56 |
| 2 | 0 | Cam7_21 | Schie5_57 |
| 5 | 3 | Cam7_21 | Spar11B_64 |
| 2 | 2 | Cam7_21 | Spar11B_67 |
| 2 | 0 | Cam7_21 | Spar11B_68 |
| 5 | 3 | Cam7_21 | Saal2_94 |
| 2 | 2 | Cam7_21 | Saal2_95 |
| 2 | 0 | Cam7_21 | Saal2_100 |
| 2 | 2 | Cam7_23 | Schie5_55 |
| 3 | 2 | Cam7_23 | Schie5_56 |
| 3 | 1 | Cam7_23 | Schie5_57 |
| 2 | 2 | Cam7_23 | Spar11B_64 |
| 3 | 1 | Cam7_23 | Spar11B_67 |
| 3 | 1 | Cam7_23 | Spar11B_68 |
| 2 | 2 | Cam7_23 | Saal2_94 |
| 3 | 1 | Cam7_23 | Saal2_95 |
| 3 | 1 | Cam7_23 | Saal2_100 |
| 0 | 0 | Cam7_30 | Schie5_55 |
| 3 | 1 | Cam7_30 | Schie5_56 |
| 3 | 2 | Cam7_30 | Schie5_57 |
| 0 | 0 | Cam7_30 | Spar11B_64 |
| 3 | 1 | Cam7_30 | Spar11B_67 |
| 3 | 3 | Cam7_30 | Spar11B_68 |
| 1 | 1 | Cam7_30 | Saal2_94 |
| 3 | 1 | Cam7_30 | Saal2_95 |
| 2 | 2 | Cam7_30 | Saal2_100 |
| 5 | 3 | Schie5_55 | Spar11B_64 |
| 2 | 2 | Schie5_55 | Spar11B_67 |
| 2 | 0 | Schie5_55 | Spar11B_68 |
| 5 | 3 | Schie5_55 | Saal2_94 |
| 2 | 2 | Schie5_55 | Saal2_95 |
| 2 | 0 | Schie5_55 | Saal2_100 |
| 2 | 2 | Schie5_56 | Spar11B_64 |
| 3 | 1 | Schie5_56 | Spar11B_67 |
| 3 | 1 | Schie5_56 | Spar11B_68 |
| 2 | 2 | Schie5_56 | Saal2_94 |
| 3 | 1 | Schie5_56 | Saal2_95 |
| 3 | 1 | Schie5_56 | Saal2_100 |
| 0 | 0 | Schie5_57 | Spar11B_64 |
| 3 | 1 | Schie5_57 | Spar11B_67 |
| 3 | 3 | Schie5_57 | Spar11B_68 |
| 1 | 1 | Schie5_57 | Saal2_94 |
| 3 | 1 | Schie5_57 | Saal2_95 |
| 2 | 2 | Schie5_57 | Saal2_100 |
| 5 | 3 | Spar11B_64 | Saal2_94 |
| 2 | 2 | Spar11B_64 | Saal2_95 |

| | | | |
|---|---|---|---|
| 2 | 0 | Spar11B_64 | Saal2_100 |
| 2 | 2 | Spar11B_67 | Saal2_94 |
| 3 | 1 | Spar11B_67 | Saal2_95 |
| 3 | 1 | Spar11B_67 | Saal2_100 |
| 1 | 1 | Spar11B_68 | Saal2_94 |
| 3 | 1 | Spar11B_68 | Saal2_95 |
| 2 | 2 | Spar11B_68 | Saal2_100 |
